# Parkinson-associated *SNCA* enhancer variants revealed by open chromatin in mouse dopamine neurons

**DOI:** 10.1101/364257

**Authors:** Sarah A. McClymont, Paul W. Hook, Alexandra I. Soto, Xylena Reed, William D. Law, Samuel J. Kerans, Eric L. Waite, Nicole J. Briceno, Joey F. Thole, Michael G. Heckman, Nancy N. Diehl, Zbigniew K. Wszolek, Cedric D. Moore, Heng Zhu, Jennifer A. Akiyama, Diane E. Dickel, Axel Visel, Len A. Pennacchio, Owen A. Ross, Michael A. Beer, Andrew S. McCallion

## Abstract

The progressive loss of midbrain (MB) dopaminergic (DA) neurons defines the motor features of Parkinson disease (PD) and modulation of risk by common variation in PD has been well established through GWAS. Anticipating that a fraction of PD-associated genetic variation mediates their effects within this neuronal population, we acquired open chromatin signatures of purified embryonic mouse MB DA neurons. Correlation with >2,300 putative enhancers assayed in mice reveals enrichment for MB cis-regulatory elements (CRE), data reinforced by transgenic analyses of six additional sequences in zebrafish and mice. One CRE, within intron 4 of the familial PD gene *SNCA*, directs reporter expression in catecholaminergic neurons of transgenic mice and zebrafish. Sequencing of this CRE in 986 PD patients and 992 controls reveals two common variants associated with elevated PD risk. To assess potential mechanisms of action, we screened >20,000 DNA interacting proteins and identify a subset whose binding is impacted by these enhancer variants. Additional genotyping across the *SNCA* locus identifies a single PD-associated haplotype, containing the minor alleles of both of the aforementioned PD-risk variants. Our work posits a model for how common variation at *SNCA* may modulate PD risk and highlights the value of cell context-dependent guided searches for functional non-coding variation.

## INTRODUCTION

Parkinson disease (PD) is a common progressive neurodegenerative disorder characterized by preferential and extensive degeneration of dopaminergic (DA) neurons in the *substantia nigra*^1,2^. This loss of midbrain (MB) DA neurons disrupts the nigrostriatal pathway and results in the movement phenotypes observed in PD. While this disorder affects approximately 1% of people over 70 years old worldwide^3^, the mechanisms underlying genetic risk of sporadic PD in the population remains largely unknown. Familial cases of PD with known pathogenic mutations are better understood but account for ≤10% of PD cases^4^.

The α-synuclein gene (*SNCA*) is commonly disrupted in familial PD through missense mutations predicted to promote misfolding^5–7^ or genomic multiplications, resulting in an over-expression paradigm^8^. The *SNCA* locus has also been shown by genome-wide association studies (GWAS) to harbour common variation modulating risk of sporadic PD, with far stronger association signals observed in comparison to all other nominated PD risk loci^9^. In the same way, common variation at over 40 additional loci have been implicated in PD^10^, but the genes modulated and causal variants responsible for elevating risk remain largely undetermined.

That most GWAS-implicated variants are non-coding^11^ is a major source of this uncertainty, obstructing the identification of: 1) the causative variant at a locus; 2) the context in which the variation is acting and; 3) the mechanism by which a variant asserts its effect on disease risk.

GWAS are inherently biologically agnostic and their exploitation of linkage disequilibrium (LD) structure frequently results in many variants being implicated at any one locus, with no one variant prioritized over those in LD. One method to prioritize non-coding variants is to examine the chromatin status at that locus^11–13^. Accessible chromatin is more likely to be functional and variants therein may impact that activity, more so than those variants residing in inaccessible chromatin. Chromatin accessibility is dynamic, often varying across cell types and developmental time. Understanding the cellular context in which variation acts is therefore critical to begin the process of prioritizing variants and querying their methods of action^11,14–16^.

Exploiting the preferential vulnerability of MB DA neurons in PD, we have prioritized DA neurons as the biological context in which a fraction of PD-associated variation likely acts. DA neurons in other brain regions, such as the forebrain (FB), provide a related substrate that is less vulnerable to loss in PD. We sought to use chromatin data from *ex vivo* populations of DA neurons to investigate the contributions of non-coding variation to PD risk. To maximize the specificity of the biological context, we generated chromatin signatures of purified mouse MB and FB DA neurons. We examined the resulting regulatory regions for their ability to direct *in vivo* reporter expression and developed a regulatory sequence vocabulary specific to DA neurons. In doing so, we identified a novel MB DA regulatory element that falls within intron 4 of *SNCA* and demonstrate its ability to direct reporter expression in catecholaminergic neurons of transgenic mice and zebrafish. Furthermore, this enhancer harbours two common variants falling in a haplotype that we determine to be associated with PD risk. We demonstrate these enhancer variants impact protein binding and we propose a model for how the variants and the haplotype at large contribute to *SNCA* regulatory control. This work illustrates the power of cell context-dependent guided searches for the identification of disease associated and functional non-coding variation.

## RESULTS

### ATAC-seq identifies open chromatin in MB and FB DA neurons

To identify regions of open chromatin in DA neurons, we performed ATAC-seq^17^ on ~50,000 fluorescent-activated cell sorting (FACS)-isolated cells (per replicate) from microdissected regions of the MB and FB of embryonic day 15.5 (E15.5) Tg(Th-EGFP)DJ76Gsat BAC transgenic mice^18^ (**Figure 1a**). This mouse line expresses EGFP under the control of the tyrosine hydroxylase (*Th*) locus, labeling catecholaminergic neurons (i.e.: DA, noradrenergic, and adrenergic neurons). To confirm capture of the corresponding catecholaminergic neurons, we performed RT-qPCR on the isolated reporter-labelled cells, establishing them to be enriched for DA neuronal markers relative to unlabelled populations from the same dissected tissues (**Supplemental Figure 1**).

**Figure 1.**
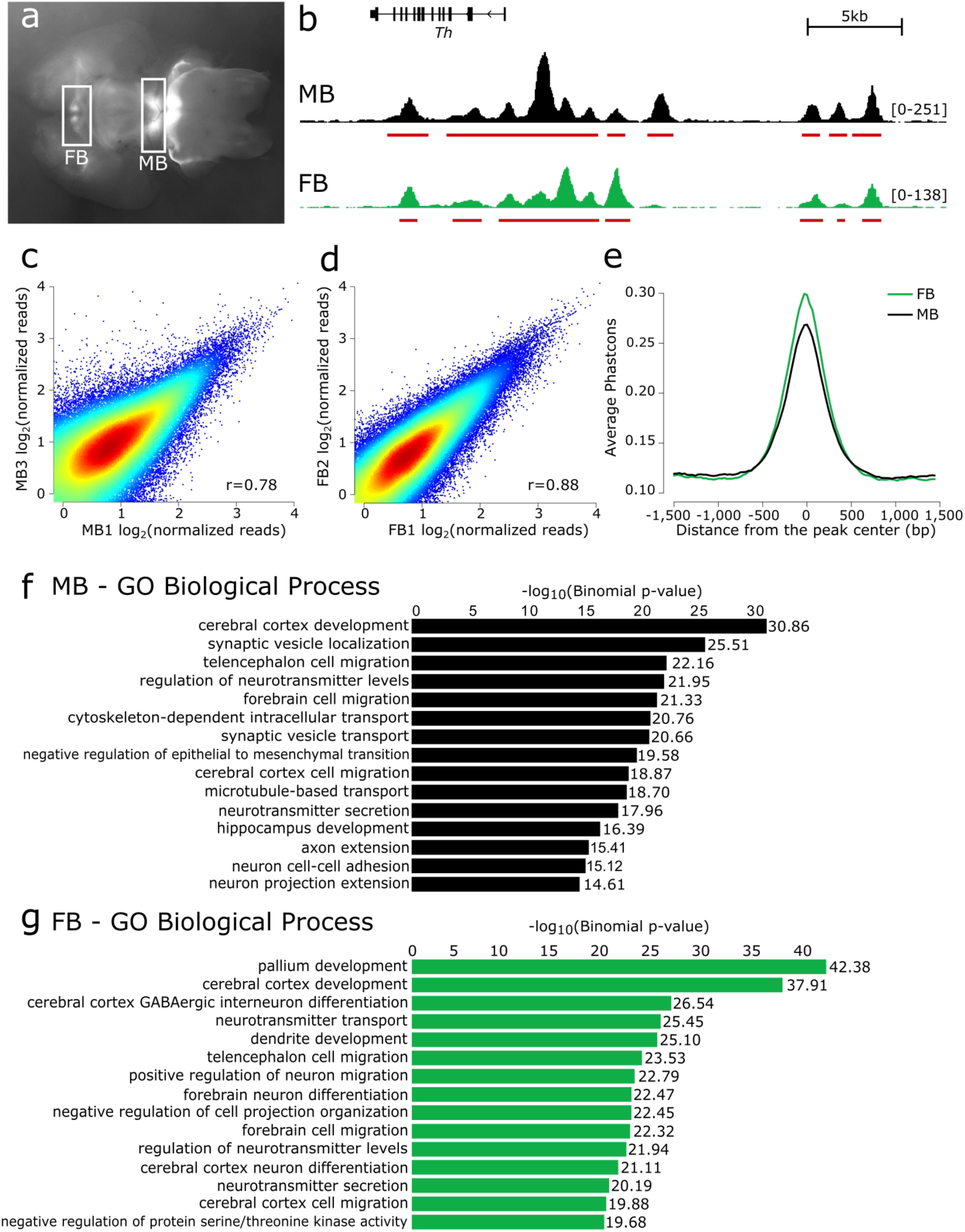
Preliminary validation of ATAC-seq catalogues generated from *ex vivo* DA neurons. (**a**) The midbrain (MB) and forebrain (FB) of E15.5 brains from Tg(Th-EGFP)DJ76Gsat mice are microdissected, dissociated, and isolated by FACS. (**b**) Read pile-up and called peaks for the MB and FB libraries at the *Th* locus. (**c, d**) Chromatin accessibility, genome-wide, is correlated between replicates. (**e**) The sequences underlying MB and FB peaks display a high degree of evolutionary sequence constraint as measured by PhastCon scores. (**f, g**) Gene ontology terms of the nearest expressed genes to all peaks in both the MB and FB reflect the neuronal origin and function of these catalogues.

To evaluate the ATAC-seq libraries, we examined the called peaks and read pile-ups with the Integrative Genomics Viewer (IGV)^19,20^ and quantified the correlation between brain regions and within replicates. A representative browser trace at the *Th* locus in both MB and FB libraries is presented in **Figure 1b**. Replicates are well correlated: MB library replicates have an average correlation of 0.72 (**Figure 1c**), and FB replicates are more correlated at r = 0.86 (**Figure 1d**). Given the robust correlation between replicates, we pooled all reads from the same brain region and called peaks on this unified set to increase our power to detect regions of open chromatin. As a result, we identified 104,217 regions of open chromatin in the MB DA neurons and 87,862 regions in the FB. MB and FB libraries are moderately well correlated (average r = 0.64; **Supplemental Figure 2**), with approximately 60% of MB peaks also represented in the FB libraries.

To assess these catalogues for characteristics of functionality, we examined the sequence constraint underlying the called regions of open chromatin, excluding peaks that overlap promoters. Promoters are typically accessible^21^ and thus, we aimed to reduce the inflation of sequence conservation due to highly conserved promoter-overlapping ATAC-seq peaks. Despite removal of these highly conserved peaks, we observed a high degree of sequence constraint underlying open chromatin peaks compared to background (**Figure 1e**). The fact that elements in these libraries of putative cis-regulatory elements (CREs) are constrained, highlights their likely functional significance.

To further examine the catalogues for biological relevance, we explored the gene ontology (GO) terms of nearby genes. While CREs are not restricted to acting solely on the nearest gene, this restriction is often used as a proxy in the absence of other information. To bolster our predictions, we also generated bulk RNA-seq data on these same populations of sorted cells and used these data to examine the GO terms of the nearest expressed gene (RPKM ≥ 1). While still imperfect, implementing this as a proxy for function results in GO terms enriched for neuronal functions in both MB and FB catalogues (**Figures 1f, g**). Thus, we establish these catalogues are enriched for putative CREs likely directing the expression of genes with key roles in neuronal biology.

### Candidate regulatory regions are capable of directing expression *in vivo*

Although our candidate CRE catalogues appear to be enriched for functional elements on the basis of sequence conservation and GO, both of these metrics are indirect surrogates for true measures of function. To more directly measure the biological relevance of the catalogues and to identify enhancers, we assessed the capability of the candidate CREs to direct expression *in vivo*.

We took advantage of the large repository of elements that have already been tested in *lacZ* reporter assays *in vivo* and catalogued in the VISTA enhancer browser^22^ (accessed September 4, 2016). Overlap between our catalogues and all 2,387 elements in the VISTA enhancer browser, which were scored for their ability to direct *lacZ* reporter expression in E11.5 mice, was quantified (**Supplemental Table 1**). Of the 1,264 elements in VISTA identified as enhancers, 786 were present in the MB catalogue and 719 were present in the FB catalogue (**Figure 2a**). Examining the overlap of the FB and MB catalogues with enhancers demonstrated to direct expression in either non-neuronal or neuronal tissues, we observed that 42-47% of enhancers reported to direct expression in non-neuronal tissues are present in the catalogues. By contrast, 71-76% of enhancers that direct expression in one or more regions of the brain overlapped the FB and MB catalogues (**Figure 2b**) confirming an abundance of brain enhancers in our catalogues. Stratifying these confirmed neuronal enhancers on the basis of their expression patterns in VISTA, we observed an abundance of MB-specific enhancers in our MB catalogue, and an abundance of FB-specific enhancers in our FB catalogue, with 77% of MB‐ and FB-specific enhancers in VISTA captured in our MB and FB catalogues, respectively (**Figure 2c**). Collectively, these data establish that our region-specific catalogues capture region-specific, active CREs with high efficiency.

**Figure 2.**
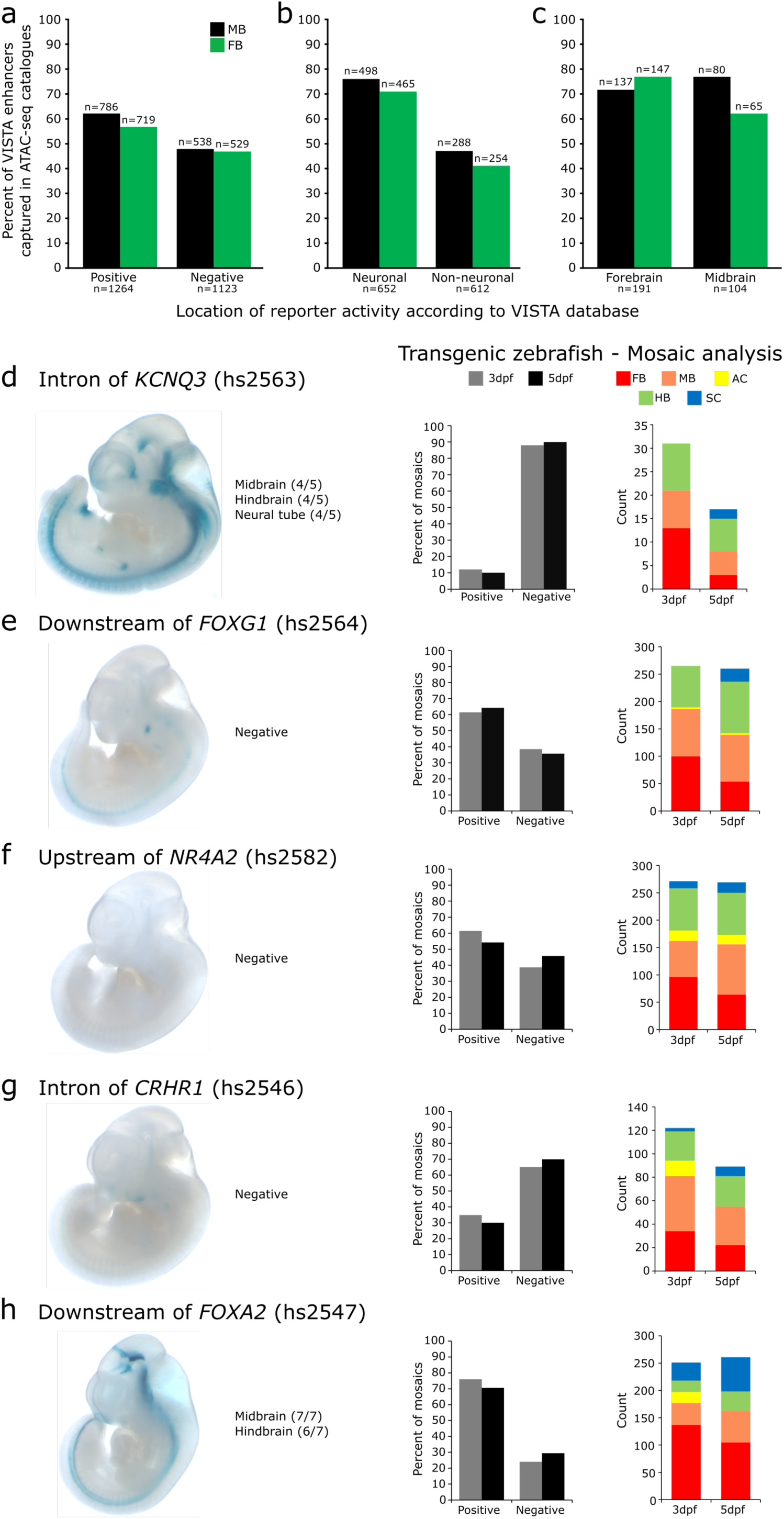
Validation of the putative CRE catalogues *in vivo*. (**a**) Of the elements annotated in VISTA as having enhancer activity, 62% and 56% of these are represented in the MB and FB catalogues, respectively (**b**) An abundance of open chromatin regions in the MB and FB catalogues overlap confirmed neuronal enhancers (≥70%). (**c**) Stratifying neuronal enhancers, MB‐ and FB-specific enhancers are enriched in our MB and FB catalogues, respectively. (**d-h**) Testing five prioritized putative CREs *in vivo* identifies five neuronal enhancers. (**d**) A putative CRE in intron 1 of *KCNQ3* directs expression in the midbrain, hindbrain, and neural tube of E11.5 lacZ reporter mice. It fails to direct expression in a transgenic zebrafish assay at either 3 or 5 days post fertilization (dpf); reporter expression present in ≤25% of mosaics. (**e, f, g**) Putative CREs downstream of *FOXG1*, upstream of *NR4A2*, and in an intron of *CRHR1* fail to direct expression in transgenic mice, however, they direct robust neuronal appropriate expression in transgenic zebrafish reporter assays (scored for expression in MB, FB, amacrine cells (AC), hindbrain (HB), spinal cord (SC)). (**h**) A putative CRE downstream of *FOXA2* directs neuronal expression in both transgenic mice and zebrafish assays. N mosaic zebrafish scored: ≥ 141 for 3dpf, ≥ 119 for 5dpf. All constructs have since been deposited in the VISTA database, under the hs numbers supplied.

To extend our assessment of the biological activity of sequences within these catalogues, we focused on an additional five candidate CREs not already tested in the VISTA browser and evaluated their ability to act as enhancers in *lacZ* reporter mice and in transgenic zebrafish TdTomato reporter assays. All five regions were represented by robust peaks in both the MB and FB catalogues (**Supplemental Figure 3**). Two regions, one in the first intron of *Kcnq3* and the other downstream of *Foxg1*, were additionally prioritized using H3K27Ac ChIP-seq from a variety of tissues from E11.5 and E15.5 embryonic mice, seeking to limit our selection to candidate enhancers predicted to have neuronal-specific activity. The remaining three candidate CREs were selected on their proximity to genes important in DA neuron biology. We selected sequences at *Foxa2* and *Nr4a2*, as both are key transcription factors (TFs) in the development and maintenance of DA neurons^23–26^. The final region, located in an intron of *Crhr1*, was selected as this locus has been implicated in PD by GWAS^9^ and our group has recently prioritized this gene as a candidate for PD risk^27^. All selected sequences were lifted over to hg19, facilitating the identification and assay of their corresponding human sequence intervals.

When tested in transgenic reporter mice at E11.5 (**Supplemental Figure 3**), two of the five regions (those near *KCNQ3* and *FOXA2*) were validated as enhancers (**Figure 2c, g**). Recognizing that a disparity exists between the developmental time at which we generated the catalogues (E15.5) and when the data was assayed (E11.5), which may compromise validation rates, we also assayed each sequence across multiple time points in zebrafish. All assayed regions except that at *KCNQ3* directed reporter expression in mosaic transgenic zebrafish (**Figures 2d, e, f, g**). All five regions displayed enhancer activity *in vivo* in neuronal tissues in one or both transgenic assays. Our transgenic animal experiments corroborate the results of the retrospective VISTA enhancer browser intersection; implying that our catalogues of candidate CREs are biologically active and enriched for sequences capable of driving neural expression *in vivo*.

### Candidate CREs are enriched for TF motifs active in DA neurons

To identify sequence modules (kmers) predicted to contribute regulatory activity of putative CREs in our catalogues, we applied the machine learning algorithm, gkm-SVM^28^. The resulting regulatory vocabularies of kmers had high predictive power (auROC_MB_ = 0.915, auROC_FB_ = 0.927). We rank ordered and collapsed related kmers to reveal motifs enriched in the putative CREs and their corresponding TFs (**Figures 3a, e, i, m**). In the MB, the four most enriched motifs correspond to Rfx1, Foxa2, Ascl2, and Nr4a2. Given the degeneracy of binding motifs within TF families, we consulted the bulk RNA-seq data for each of the implicated TF families and examined the relative expression levels to prioritize which TFs are most likely producing the observed motif enrichments (**Figures 3b, f, j, n**). While no member of the Rfx family has been canonically associated with MB DA neurons, we anticipate Rfx3 and Rfx7, as the two highest expressed Rfx genes, to likely be active in MB DA neurons and driving this motif enrichment (**Figure 3b**). Foxa1, and more specifically, Foxa2 are both known to DA neuron biology^23,29^ and both are highly expressed in the MB DA neurons (**Figure 3f**). Regarding enrichment for the Ascl family, Ascl1 is known to be involved in DA neuron biogenesis^30^ and is more highly expressed than any other TF in the family (**Figure 3j**). Finally, Nr4a2 is both canonically associated with DA neurons and required for their development^26^; we observe it to be highly expressed in MB DA neurons (**Figure 3n**). Examining the sequences underlying the CRE catalogues, we identified TF families known and unknown to DA neuron biology and further refined the TF associations using expression data.

**Figure 3.**
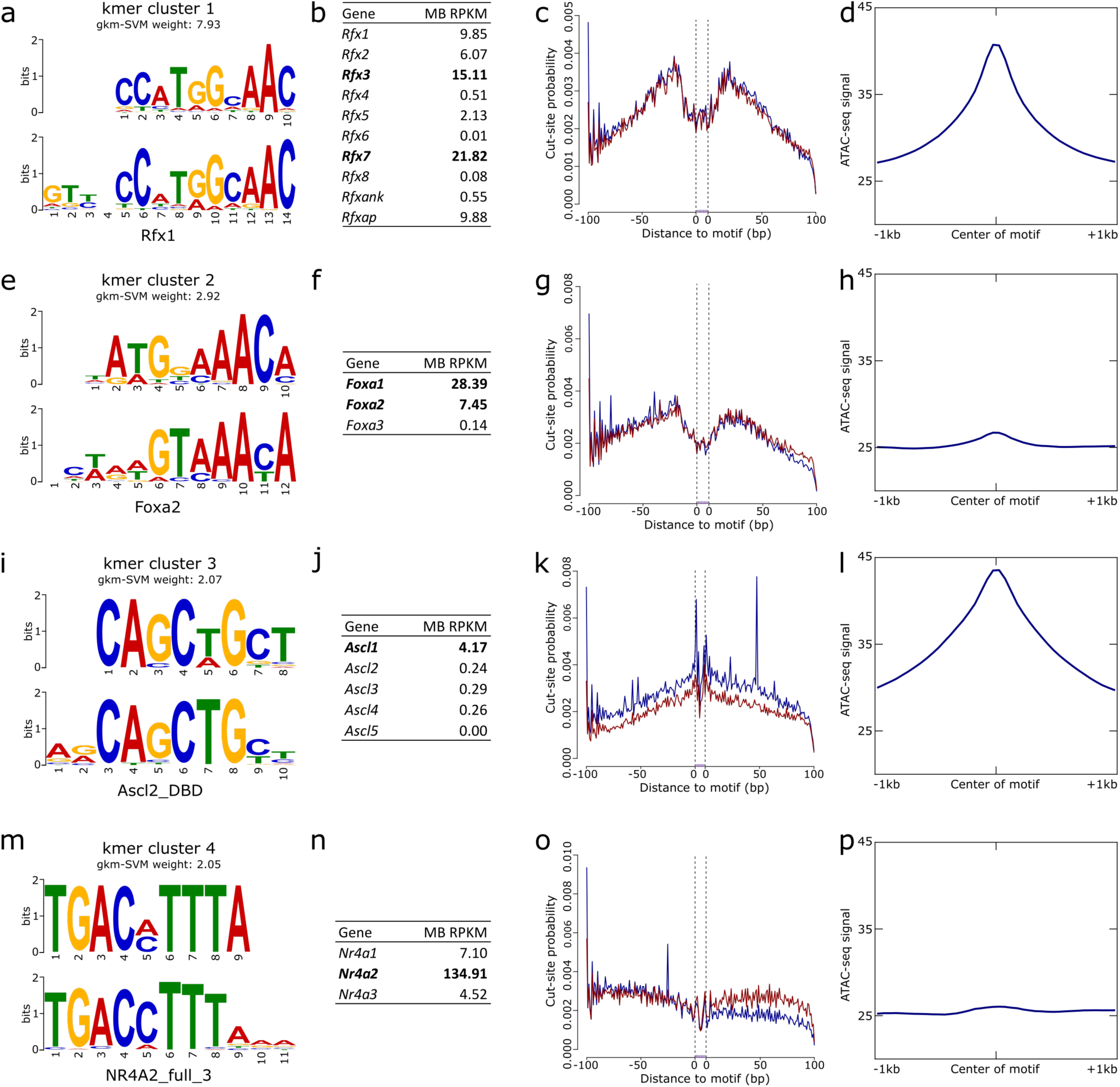
Identification of transcription factors (TFs) important to DA neurons. (**a**) The kmer predicted to have the greatest regulatory potential underlying MB ATAC-seq peaks corresponds to the Rfx family of TFs. (**b**) RNA-seq quantification in these same cells indicates this enrichment is likely due to Rfx3 or Rfx7 activity. Examining the ATAC-seq signal over predicted binding sites reveals a robust TF footprint (**c**) and a general enrichment of reads overlapping Rfx sites genome-wide (**d**). (**e-h**) Similarly, a kmer corresponding to the TFs Foxa1/2 have similar evidence for their activity. (**i-j**) The third ranked motif likely corresponds to Ascl1, and while it fails to leave a robust TF footprint (**k**), there is clear enrichment of ATAC-seq signal overlapping genome-wide predicted Ascl1 binding sites (**l**). (**m-n**) Nr4a2, canonically associated with DA neuron biology, is identified as a highly expressed TF likely contributing to the regulatory potential of the putative CREs however, it fails to leave a TF footprint in the cut-site patterns around predicted motif sites (**o**) and is only mildly enriched for ATAC-seq reads over its predicted binding sites (**p**).

We also examined the qualities that differentiate MB CREs from FB CREs by examining the sequences underlying MB-specific and FB-specific regions. We developed a vocabulary that discriminates MB and FB regions with high predictive power (auROC = 0.926) and identified kmers enriched in MB-specific peaks where the top corresponding TFs are Foxa1/2 and Nr4a2 (**Supplemental Figure 4**). We confirmed this MB bias by again considering the bulk RNA-seq for these genes. As expected, these TFs are more highly expressed in the MB where *Nr4a2* is present at 12-fold higher levels in the MB (135 RPKM in the MB vs 11 RPKM in the FB) and *Foxa1/2* are not expressed in the FB, but are present in the MB (*Foxa1*: 28 RPKM, *Foxa2*: 7 RPKM). Not only do we identify Foxa1/2 and Nr4a2 as more active in MB DA neurons than in the FB, we did so solely by comparing their role in the vocabulary of MB-specific candidate CREs versus FB-specific CREs.

In a parallel strategy to identify TFs actively engaging the DNA in MB DA neurons, we performed TF footprinting in a single deeply sequenced MB ATAC-seq library. Doing so, we confirm that two of the TFs prioritized by gkm-SVM leave robust footprints. The motif corresponding to Rfx binding results in a dearth of cuts directly over predicted binding sites (**Figure 3c**). The same can be seen to a lesser extent for the motif corresponding to Foxa1/2 (**Figure 3g**). By contrast, motifs corresponding to Ascl1 or Nr4a2 fail to leave a robust mark on the chromatin availability (**Figures 3k, o**). These footprinting data substantiate the claim that the Rfx family of TFs and Foxa1/2 are active in MB DA neuron CREs.

We confirmed that these sequences are indeed enriched in the catalogues by examining the pileup of reads overlapping all genome-wide predicted motif binding sites for each motif identified by gkm-SVM. We see an abundance of reads over predicted binding sites of all four motifs (**Figures 3d, h, l, p**), with the strongest enrichment overlapping Rfx and Ascl1 motif sites (**Figures 3d, l**). Despite the less robust footprint generated at the Ascl1, this TF clearly underlies a larger than expected proportion of CREs in the MB catalogue. The integration of a support vector machine learning algorithm as applied to the sequences underlying open chromatin regions with footprinting analysis in the same chromatin substrate powerfully identifies TFs that are important for DA neuron biology and suggests the Rfx family of TFs, Foxa1/2, Ascl1, and Nr4a2 are actively influencing gene expression in the MB DA neurons.

### A candidate CRE in intron 4 of SNCA is associated with PD risk

Having established the biological robustness of the CRE catalogue, we moved to exploit these data to investigate how non-coding variation therein may be contributing to PD risk; given α-synuclein’s established role in PD pathogenesis, we prioritized this locus for investigation. We first noted that *Snca* expression differs significantly between the MB and FB DA neurons in our bulk RNA-seq (**Figure 4b**). Examining the chromatin accessibility at the *Snca* locus, the MB and FB are largely the same with the exception of one robust peak in intron 4 (mm9: chr6:60,742,503-60,744,726) that is present in the MB and completely absent in the FB (**Figure 4a**). DNase hypersensitivity site (DHS) linkage^21,31^ suggests that this putative CRE interacts with the *SNCA* promoter. Given the MB-specificity of this putative CRE and indications that it interacts with the *SNCA* promoter, we anticipated this region to be a driving force behind the MB-specific expression of *Snca*.

**Figure 4.**
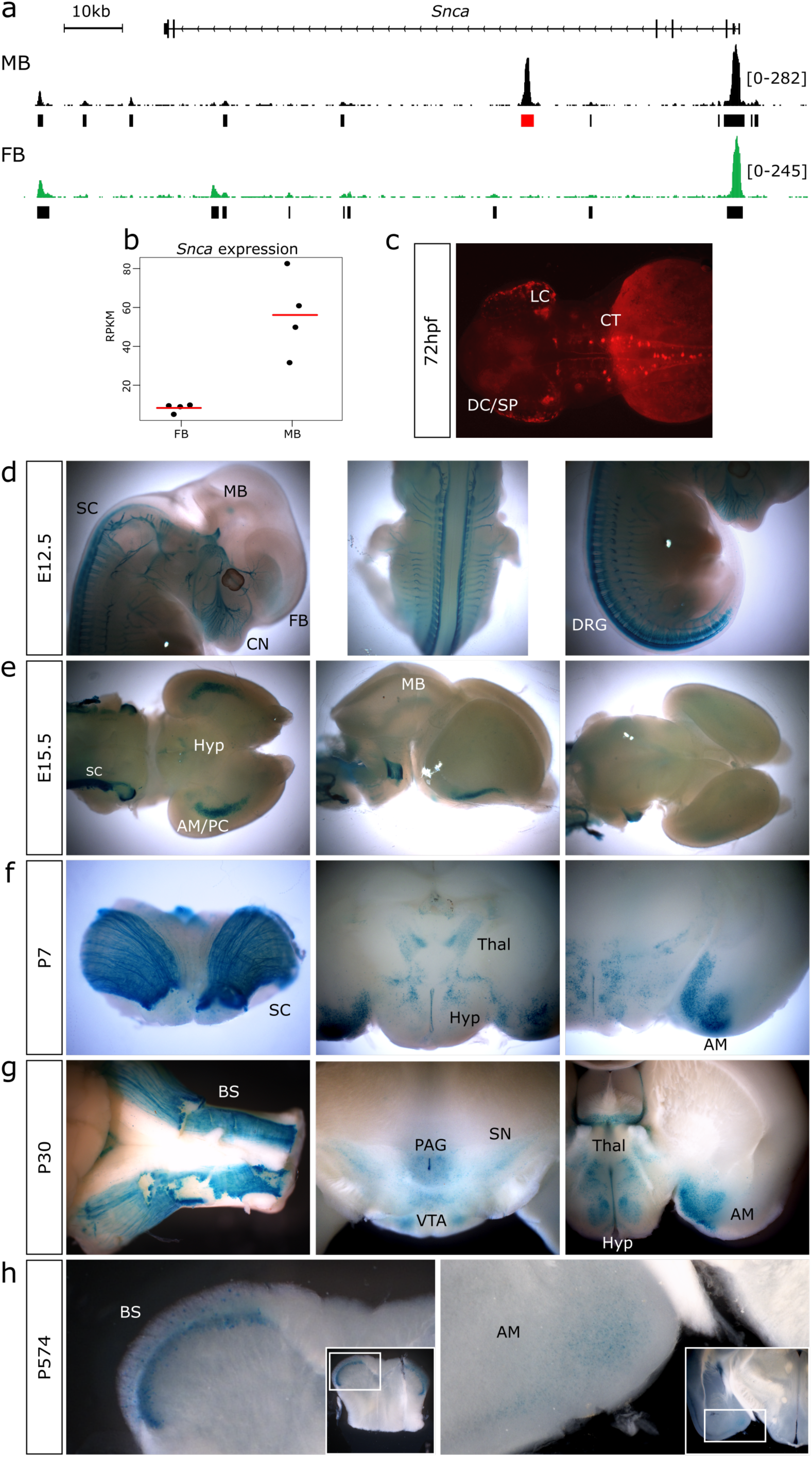
A MB-specific enhancer directs expression in catecholaminergic populations of neurons known to Parkinson disease biology. (**a**) IGV track indicating the location of the MB-specific region of open chromatin, located in intron 4 of *Snca*. (**b**) *Snca* is differentially expressed between the MB and FB DA neurons. Red bar is the mean expression of the four replicates (black dots). (**c**) At 72 hours post fertilization (hpf), stable transgenic zebrafish reporter assays indicate this putative CRE is capable of directing reporter expression in key catecholaminergic neuronal populations, including the locus coeruleus (LC), the catecholaminergic tract (CT) of the hindbrain, and the diencephalic cluster (DC) with projections to the subpallium (SP). (**d-g**) Further studies in *lacZ* reporter assays in embryonic (**E**) and post-natal (**P**) mice indicate dynamic enhancer usage across developmental time. (**d**) This enhancer directs expression throughout the MB, FB, dorsal root ganglia (DRG), sympathetic chain (SC), and cranial nerves (CN) of E12.5 mice. (**e**) By E15.5, reporter expression is observed in the amygdala and/or piriform cortex (AM/PC), sympathetic chain, MB, and hypothalamus (Hyp). (**f**) Patterns of reporter expression at P7 reflect those seen at E15.5. (**g**) Reporter activity is observed at P30 in the amygdala, hypothalamus and thalamus (Thal), brain stem (BS), *substantia nigra* (SN), ventral tegmental area (VTA), and the periaqueductal grey area (PAG). (**h**) In aged mice (P574), reporter expression is detected robustly in the brain stem and faintly in the amygdala.

To test this hypothesis, we assayed whether the central portion of this putative CRE, when lifted over to hg19 (chr4:90,721,063-90,722,122), is capable of directing appropriate reporter expression in transgenic zebrafish and mouse reporter assays. Stable transgenesis of zebrafish indicates that this CRE directs reporter expression at 72 hours post fertilization in the locus coeruleus, a key population of catecholaminergic neurons preferentially degenerated in PD^32^, and along the catecholaminergic tract through the hindbrain, which is largely composed of DA neurons^33^ (**Figure 4c**). Additionally, we observe reporter expression throughout the diencephalic catecholaminergic cluster with projections to the subpallium, which is analogous to mammalian dopaminergic projections from the ventral midbrain to the striatum^34^. Reporter expression in these transgenic zebrafish is largely consistent with an enhancer active in catecholaminergic populations.

To further evaluate this CRE in a mammalian system, we generated *lacZ* reporter mice and examined reporter activity across developmental time. Whole mount E12.5 reporter mice indicate this enhancer directs exquisitely restricted expression in Th+ populations, including the dorsal root ganglia, extending into the sympathetic chain, and throughout the cranial nerves (particularly the trigeminal). Additional diffuse staining is noted throughout the MB and FB (**Figure 4d**). Specifically examining the brains of *lacZ* animals at E15.5, reporter expression is identified in the MB and hypothalamus, with strong expression through the amygdala/piriform cortex and along the anterior portion of the sympathetic chain (**Figure 4e**); similar reporter patterns are seen at P7 (**Figure 4f**). At P30, we detect reporter activity in the amygdala, hypothalamus, thalamus, periaqueductal grey area, brain stem, and importantly, in the *substantia nigra* and ventral tegmental area (**Figure 4g**). By contrast, in aged *lacZ* reporter mice (574 days old, ~19 months), we only detect strong reporter expression in the brain stem and observe weak reporter expression in the amygdala (**Figure 4h**). Collectively, the regions in which we detect reporter activity reflect those compromised in PD; Lewy bodies (aggregates of α-synuclein) have been detected in the locus coeruleus, sympathetic chain, amygdala, hypothalamus, ventral tegmental area, periaqueductal grey area of PD patients^35–39^, and critically the preferential degradation of the *substantia nigra* is the pathological hallmark of PD progression^2^. This enhancer directs region-specific appropriate expression throughout development in key locations concordant with SNCA activity in PD pathogenesis.

Following confirmation of this CRE’s regulatory activity in brain regions associated with PD, we next inspected this sequence for PD-associated variation. We sequenced across this interval in 986 PD patients and 992 controls and identified 14 variants (**Supplemental Table 2**), 4 of which were common and present in both cases and controls with a minor allele frequency greater than 5%. Of these, two tightly linked variants (r^2^ = 0.934; **Supplemental Table 3**), rs2737024 (OR = 1.25, 95% CI = 1.09-1.44, *p*-value = 0.002) and rs2583959 (OR = 1.22, 95% CI = 1.06-1.40, *p*-value = 0.005), were significantly associated with PD (**Table 1**). These data support a role for variation within the enhancer in conferring PD risk.

**Table 1:** Two tightly linked SNPs within the enhancer are significantly associated with PD risk

| Variant | MA | MAF in PD cases<br>(N=986) | MAF in controls<br>(N=992) | Association with PD |  |
| --- | --- | --- | --- | --- | --- |
|  |  |  |  | OR (95% CI) | p-value |
| rs7684892 | A | 0.069 | 0.065 | 0.93 (0.72-1.20) | 0.562 |
| rs17016188 | C | 0.082 | 0.061 | 1.35 (1.04-1.75) | 0.023 |
| rs2583959 | G | 0.317 | 0.271 | 1.22 (1.06-1.40) | 0.005 * |
| rs2737024 | G | 0.319 | 0.270 | 1.25 (1.09-1.44) | 0.002 * |

MA=minor allele; MAF=minor allele frequency; OR=odds ratio; CI=confidence interval Only variants with MAF > 0.05 were considered ORs, 95% CIs, and p-values result from additive logistic regression models adjusted for age at blood draw and sex p-values ≤ 0.0125 were considered as statistically significant after applying a Bonferroni correction for multiple testing (\*)

To assess how these variants may impact enhancer function and thus PD risk, we assayed differential protein binding at these variants for >20,000 proteins^40^. In doing so, we identify five proteins whose binding is robustly impacted by these implicated variants: NOVA1, APOBEC3C, PEG10, SNRPA, and CHMP5 (**Figure 5a, b, c**). Of these, all are expressed at appreciable levels in both MB and FB DA neurons (**Figure 5d**), excluding APOBEC3C (RPKM ≤ 1). Of the remaining four proteins, three (PEG10, SNRPA, and CHMP5) demonstrate an increased binding affinity for the minor risk allele over the major allele; this direction of effect is consistent with the over-expression paradigm by which *SNCA* confers PD risk^8^. Interestingly, CHMP5 is the sole protein we identify whose binding affinity is impacted by variant rs2583959, and our group has recently implicated one of its family members, CHMP7, in conferring PD risk^27^, perhaps indicating a role for this family of proteins in PD. Although no single protein stands out, the increased affinity for the risk alleles of the identified enhancer variants by proteins expressed in DA neurons is consistent with a potential mechanistic contribution to SNCA expression and therefore, PD risk.

**Figure 5.**
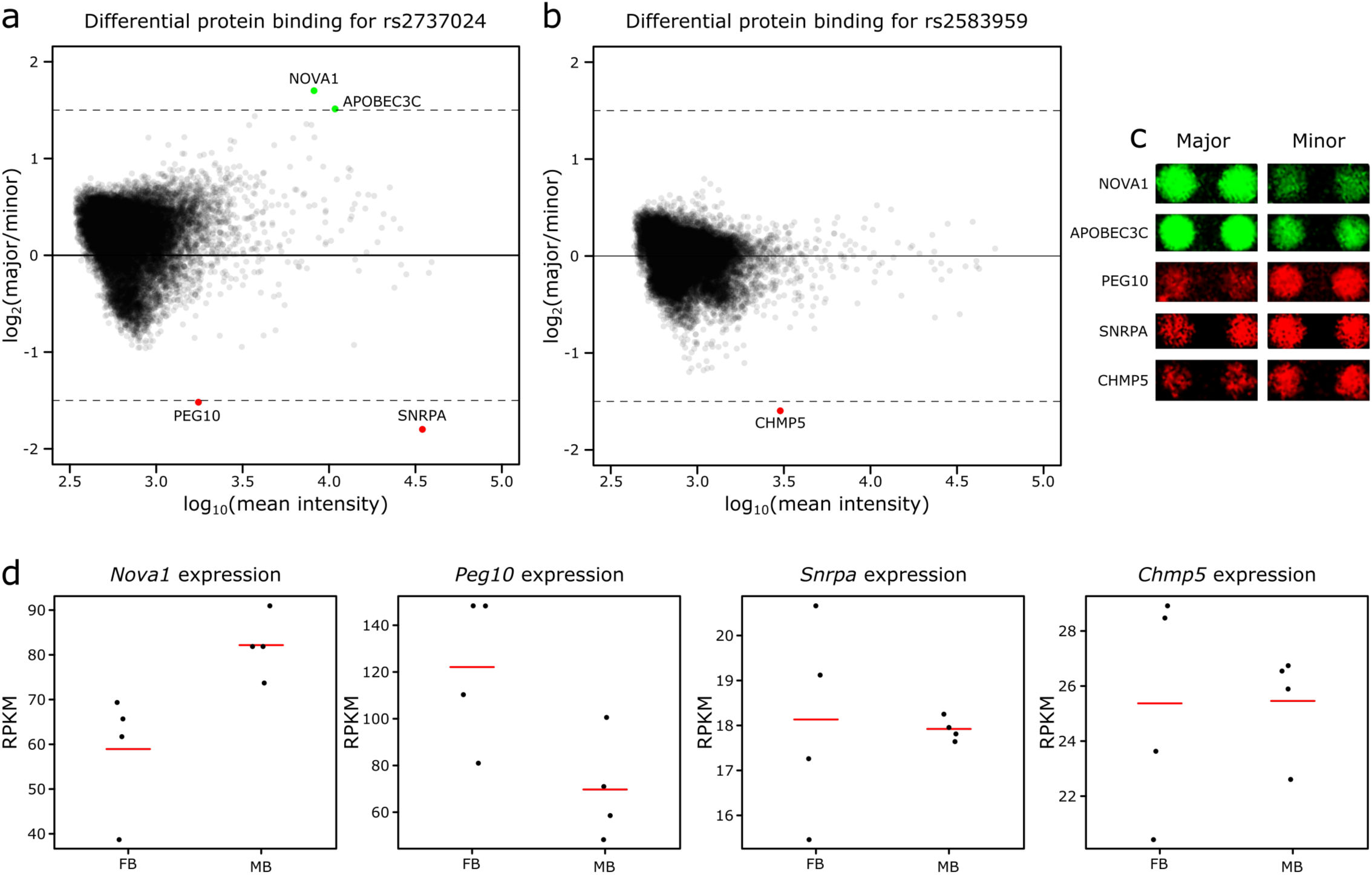
Identification of proteins whose binding is impacted by the implicated PD-risk SNPs. (**a, b**) MA plots for both rs2737024 and rs2583959 indicating the magnitude of the effect of the minor and major allele on binding. Cut-off for differential binding: log_2_(major/minor) ≥ 1.5 or ≤ −1.5. (a) NOVA1 and APOBEC3C (green circles) bind at rs2737024 with greater affinity for the major allele, while PEG10 and SNRPA (red circles) have a greater affinity for the minor allele. (**b**) CHMP5 (red circle) has a greater affinity for the minor allele of rs2583959. (**c**) Representative images of the protein binding for each of the differentially bound proteins. (**d**) Expression analysis in the MB and FB DA neurons for each of the differentially bound proteins indicate *Nova1*, *Peg10*, *Snrpa*, and *Chmp5* to be highly expressed in these populations, while none of the *Apobec* family member genes are expressed (RPKMs ≤ 1, data not shown). Red bar is the mean expression of the four replicates (black dots).

Finally, we set out to refine the haplotype structure and understand how this identified variation may be interacting with other variants at this locus. A panel of common variants had previously been genotyped across *SNCA* and PD-associated haplotypes were identified^41^. After genotyping our patients and controls for a subset of this panel of variants in addition to all enhancer-associated variants identified by sequencing (**Supplemental Table 4**), we identified a single haplotype that was significantly associated with PD (p-value = 0.003), with a higher observed frequency in PD patients (28.3%) compared to controls (23.4%; **Table 2**). This haplotype implicates some of the same variants as in Guella *et al*.^41^ (rs356220, rs737029) but also implicates rs356225 and rs356168, and the two enhancer-associated variants. Collectively, these data identify a catecholaminergic enhancer harbouring common variation that is part of a larger haplotype associated with PD risk.

**Table 2:**
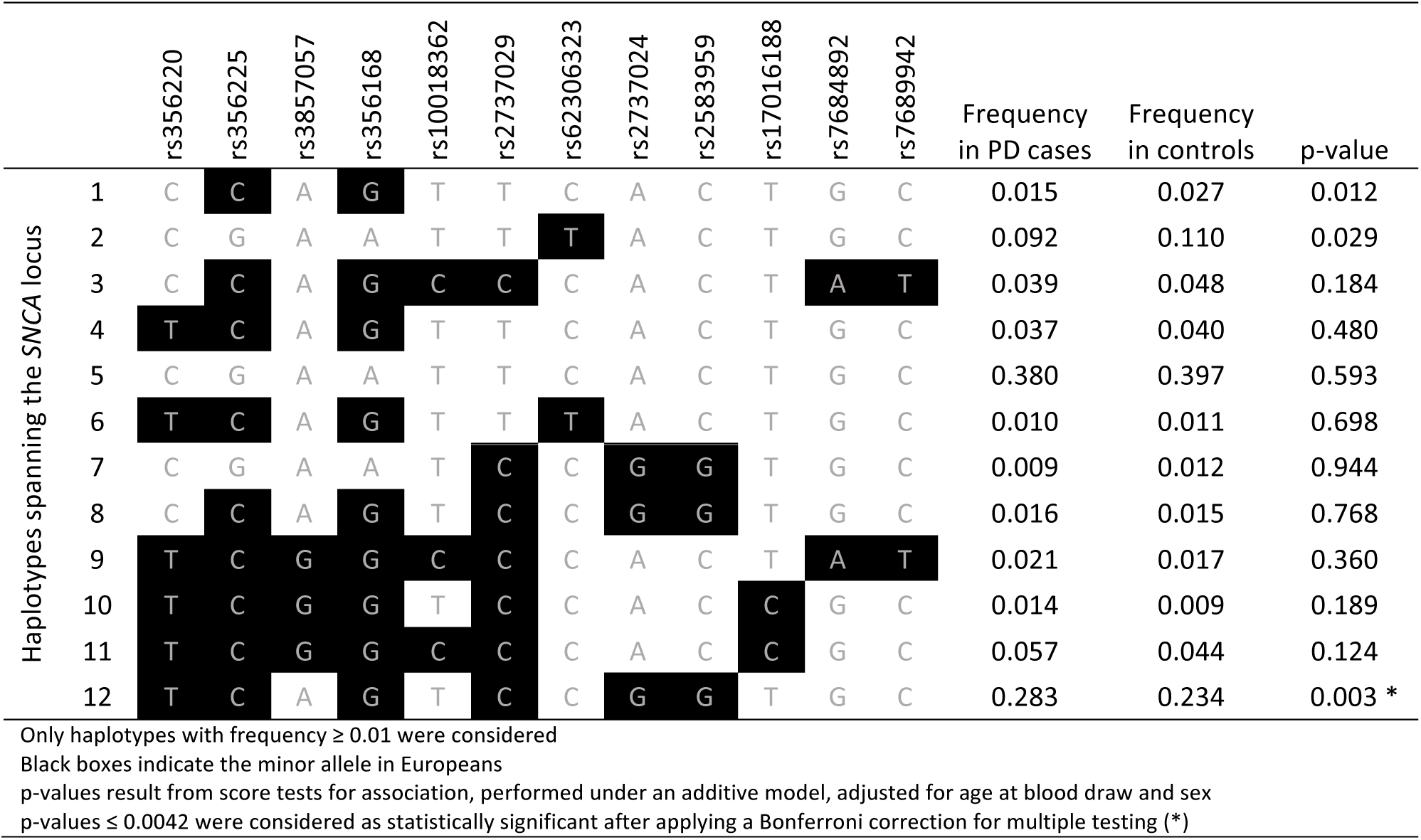
A single haplotype, containing the minor alleles of the implicated SNPs, is significantly associated with PD risk

## DISCUSSION

The identification and prioritization of biologically pertinent non-coding variation associated with disease remains challenging. Recent studies by our and other groups have emphasized the importance of cellular context in the identification of sequences harbouring biologically pertinent variation and the genes they regulate. To this end, we used chromatin signatures from *ex vivo* isolated DA neurons to reveal biologically active sequences that harbour non-coding variation contributing to PD risk. We generated robust CRE catalogues for both MB and FB DA neurons, confirmed their capacity to act as enhancers, identified motifs that confer their regulatory potential, and notably, identified two variants located within a MB-specific enhancer that are associated with an increase in PD risk.

In contrast to strategies predicated solely on dissection of post-mortem tissues or on the differentiation of cultured cells, we leveraged the use of transgenic reporter mice to specifically isolate Th-expressing neurons from discrete neuroanatomical (FB and MB) domains. While our approach assays a more refined population of DA neurons than would be achieved via gross dissection, recent single-cell RNA-seq analyses of these same cells make clear that even within these highly restricted MB and FB populations there exist two primary cellular phenotypes^27^. The “homogenous” MB and FB populations each are comprised of an immature neuroblast population and a more mature, domain specific, post-mitotic population of DA neurons. As such, our CRE catalogues capture the chromatin accessibility from both of these states. These catalogues are demonstrably biologically relevant for our purposes, but future studies requiring even greater homogeneity may wish to consider single-cell ATAC-seq to refine these domains further^42^.

In our *in silico* validation of the catalogues, we established them to be enriched for both sequence constraint and biological relevance in a manner consistent with function and their FB/MB origin. Furthermore, these sequences are frequently domain appropriate enhancers, with each catalogue capturing a large fraction (77%) of previously validated MB and FB enhancers. Although an abundance of regions are shown to direct neuronal expression compared to those annotated as negative or non-neuronal, it is interesting to note that almost half of the sequences previously documented not to direct expression *in vivo* are also represented in one or both of our catalogues.

Given the frequently dynamic nature of CRE activity, this overlap with negative regions likely results from temporal differences in these assays. Our data indicates these regions are accessible at E15.5 but the *lacZ* reporter assays were carried out at E11.5; regions that have been annotated as negative at E11.5 may be active at later time points and, as such, appear in our catalogues. As we moved from these unbiased functional comparisons to more highly selected ones, the potential impact of temporal differences became more pronounced. In mouse transgenic reporter assays, two of five assayed putative CREs direct detectable expression of *lacZ* in neuronal populations. Consistent with the temporally dynamic nature of CREs, when these same regions are tested in zebrafish across multiple developmental time points, we observe four of the five sequences to act as neuronal enhancers.

In examining the sequence composition underlying the ATAC-seq peaks, we illuminate powerful vocabularies for both FB and MB DA neuron transcriptional regulatory control. Machine learning using gkm-SVM prioritizes four transcription factor families (Rfx, Foxa1/2, Nr4a2, Ascl1/2) as those conveying significant regulatory potential in the CRE catalogues. Of these, the Rfx family had not previously been implicated in DA neuron biology. Although several of the Rfx family members have been annotated as having expression in the cerebellum or fetal brain^43^, a role specifically in MB DA neurons has not previously been appreciated. By contrast, Nr4a2 is canonically associated with MB DA neurons^25,26^, is highly expressed in this population (139 RPKM), and was prioritized as a TF conferring regulatory potential in these cells; however, TF footprinting fails to provide evidence supporting its activity. We postulate that this lack of footprint may reflect the transient DNA binding dynamics of Nr4a2. Transcription factors with short DNA residence times often fail to reveal footprints, and nuclear receptors, such as Nr4a2, have markedly transient DNA interactions^44^.

Taken collectively, these data establish a robust biological platform in which PD-associated variation can be evaluated. To this end, an obvious candidate to interrogate was an apparent MB-specific open chromatin domain within intron 4 of the known PD-associated gene, *SNCA*. We assayed the activity of this putative CRE in zebrafish and across the life course of mice and found it to be active in key catecholaminergic structures injured in PD (e.g.: the *substantia nigra* and locus coeruleus), from mid-gestation until at least P30. Thereafter, the utilization of this enhancer in the brain is diminished and by late life appears restricted to the brainstem and amygdala. By the time of clinical presentation, PD patients have already lost a significant proportion (≥30%) of their nigral DA neurons^2,45^; the observed biology of this CRE is consistent with a progressive pathogenic influence acting early in life, rendering these populations preferentially vulnerable to loss over an extensive period of time.

Sequencing this interval in PD cases and controls revealed two common variants (rs2737024 and rs2583959) therein, individually associated with an increased risk of PD. Testing these variants for their effect on protein binding, we identify five proteins whose binding is affected, three of which, PEG10, SNRPA, and CHMP5, display greater affinity for the risk allele. Furthermore, we identify a larger haplotype containing these variants, also significantly associated with PD risk. While none of the other SNPs in this haplotype overlap with CREs identified in the DA neuron catalogues, variant rs356168 has significant functional evidence of its activity and contribution to PD risk^46^. The same DHS correlation analysis^21,31^ that suggests an interaction between the *SNCA* promoter and our identified CRE, also suggests an interaction between the SNCA promoter and the rs356168 variant. Additionally, ChIA-PET data^31,47^ indicates that sequence encompassing this variant may interact with our enhancer, suggesting a potential co-operative mode of action; a paradigm recently proposed by Gupta and colleagues^48^ at the *EDN1* locus. We propose that the variants within the enhancer, independently or in concert with other variation within the identified haplotype, may act throughout the lifespan to render key populations of catecholaminergic neurons vulnerable, thus increasing PD risk in individuals harbouring this variation.

This work emphasizes the value of biologically informed, cell context-dependent guided searches for the identification of disease associated and functional non-coding variation. Given the extent of non-coding GWAS-identified variation, the need for strategies to prioritize variants for functional follow-up is greater than ever. Here, we generate chromatin accessibility data from purified populations of DA neurons to generate catalogues of putative CREs. We have demonstrated how these data can be used to reveal non-coding variation contributing to PD risk; focusing on a single region of open chromatin at the *SNCA* locus, we uncover PD-associated variation therein and propose a model through which this sequence can contribute to normal DA neuronal biology and PD risk. There remains a plethora of information still to be explored in these catalogues, either through further single locus investigations or through massively parallel assays. For example, our MB DA neuron CRE catalogue overlaps SNPs at 10 of 26 (38%) PD-associated loci^9^, all of which can be investigated further for their mechanisms by which they impact PD risk. Our work establishes a powerful paradigm, leveraging transgenic model systems to systematically generate cell type specific chromatin accessibility data and reveal disease-associated variation, in a manner that can be progressively guided by improved biological understanding.

## METHODS

### Animal husbandry

Tg(Th-EGFP)DJ76Gsat mice (Th-EGFP) were generated by the GENSAT Project and purchased through the Mutant Mouse Resource and Research Centers Repository. Colony maintenance matings were between hemizygous male Th-EGFP mice and female Swiss Webster (SW) mice, obtained from Charles River Laboratories. This same mating scheme was used to establish timed matings, generating litters for assay; day on which vaginal plug is observed, E0.5. Adult AB zebrafish lines were maintained in system water according to standard methods^49^. All work involving mice and zebrafish (husbandry, colony maintenance, procedures, and euthanasia) were reviewed and pre-approved by the institutional care and use committee.

### Neural dissociation and FACS

Pregnant SW mice were euthanized at E15.5 and the embryos were removed and immediately placed in chilled Eagle’s Minimum Essential Media (EMEM) on ice. Embryos were decapitated and brains were removed into Hank’s Balanced Salt Solution without Mg^2+^ and Ca^2+^ (HBSS w/o) on ice. Under a fluorescent microscope, EGFP+ brains were identified and microdissected to yield the desired forebrain (FB) and midbrain (MB) regions desired. Microdissected regions were placed in fresh HBSS w/o on ice, and pooled per litter for dissociation.

Pooled brain regions were dissociated using the Papain Dissociation System (Worthington Biochemical Corporation). The tissue was dissociated in the papain solution for 30 minutes at 37°C, with gentle trituration every 10 minutes using a sterile Pasteur pipette. Following dissociation, cells were passed through a 40μm cell strainer into a 50mL conical, centrifuged for 5 minutes at 300g, resuspended in albumin-inhibitor solution containing DNase, applied to a discontinuous density gradient, and centrifuged for 6 minutes at 70g. The resulting cell pellet was resuspended in HBSS with Mg^2+^ and Ca^2+^ and submitted to FACS. Aliquots of 50,000 EGFP+ cells were sorted directly into 300μL HBSS with Mg^2+^ and Ca^2+^ with 10% FBS for ATAC-seq. Aliquots containing ≥50,000 EGFP+ cells were sorted into kit-provided lysis buffer for RNA-seq. This procedure was repeated such that a single aliquot of cells from each region per litter were submitted to either ATAC-seq or bulk RNA-seq three times over for each region.

### ATAC-seq library preparation and quantification

ATAC-seq library preparation generally follows the steps as set out in the original ATAC-seq paper^17^ with minor modifications. Aliquots of 50,000 EGFP+ cells were centrifuged for 5 minutes at 4°C and 500g, washed with 50μL of chilled PBS and centrifuged again for 5 minutes at 4°C and 500g. The cell pellet was resuspended in lysis buffer, as set out in the protocol, and cells were left to lyse for 5 minutes at 4°C before being centrifuged for 10 minutes at 4°C at 500g. The resulting nuclei pellet was transposed, as written, using the transposase from the Nextera DNA Library Preparation Kit. Following transposition, DNA was purified with the MinElute Reaction Clean-up Kit (Qiagen) and eluted in 10μL elution buffer.

Libraries were amplified according to the original ATAC-seq protocol^17^. The qPCR surveillance steps were modified such that the additional number of cycles of amplification were calculated as ¼ maximum intensity, so as to limit PCR duplication rates in the final libraries. Amplified libraries were purified with Ampure XP beads (Beckman Coulter) following the Nextera DNA Library Prep Protocol Guide. Libraries were quantified using the Qubit dsDNA High Sensitivity Assay (Invitrogen) in combination with the Agilent 2100 Bioanalyzer using the High Sensitivity DNA Assay (Agilent).

### ATAC-seq sequencing, alignment, and peak calling

Individual ATAC-seq libraries were sequenced on the Illumina MiSeq to a minimum depth of 20 million, 2×75bp reads per library. A single MB ATAC-seq library was sequenced on the Illumina HiSeq in Rapid Run mode with 2×100bp reads, to a depth of ≥350 million paired-end reads.

Quality of sequencing was evaluated using FastQC (v0.11.2; https://www.bioinformatics.babraham.ac.uk/projects/fastqc/). Reads were aligned to mm9 using Bowtie2^50^ (v2.2.5), under‐‐local mode. Reads aligning to the mitochondrial genome, unknown and random chromosomes, and PCR duplicates were removed prior to peak calling (SAMtools^51^). Peaks were called on individual libraries and on a concatenated file combining all MB and all FB libraries (“Joint”) using MACS2^52^ (v2.1.1.20160309) “callpeak” with options: ‐‐nomodel ‐‐nolambda ‐B ‐f BAMPE ‐‐gsize mm ‐‐keep-dup all. Peaks overlapping blacklisted regions called by ENCODE and in the original ATAC-seq paper were removed^17,47^.

### RNA-seq library preparation and quantification

Total RNA was extracted using the Purelink RNA Micro Kit (Invitrogen). Following FACS isolation into kit-provided lysis buffer, samples were homogenized and RNA extraction proceeded using manufacturer’s recommendations. Total RNA integrity was determined using the RNA Pico Kit (Agilent). RNA samples were sent to the Sidney Kimmel Comprehensive Cancer Center Next Generation Sequencing Core at Johns Hopkins for library preparation, using the Ovation RNA-Seq System V2 (Nugen), and sequencing.

### RNA-seq sequencing, alignment, and transcript quantification

Libraries were pooled and sequenced on Illumina’s HiSeq 2500 in Rapid Run mode with 2×100bp reads to an average depth of >90 million reads per library. Quality of sequencing was evaluated using FastQC. FASTQ files were aligned to mm9 using HISAT2^53^ (v2.0.1-beta) with ‐‐dta specified.

Aligned reads from individual samples were quantified against a reference transcriptome using the Rsubread package^54–56^ (v1.22.3) function “featureCounts” with the following options: isPairedEnd = TRUE, requireBothEndsMapped = TRUE, isGTFAnnotationFile = TRUE, useMetaFeature = TRUE. The GENCODE vM9 GTF was downloaded^57^ (date: March 30, 2016) and lifted over from the mm10 to the mm9 genome using CrossMap (v0.2.2) with default parameters^58^. This was used for quantification, in which gene-level raw counts were converted to RPKM values and means for each region were calculated.

### cDNA synthesis and RT-qPCR for DA neuron markers

RNA was extracted using the RNeasy Mini Kit (Qiagen), after sorting 50,000 cells directly into Buffer RLT. Aliquots of 50,000 non-fluorescing cells were also collected and processed in parallel. 100ng of each RNA sample was submitted to first strand cDNA synthesis using the SuperScript III First-Strand Synthesis System for RT-PCR (Invitrogen), following the Oligo(dT) method.

Primers (**Supplementary Table 5**) were designed using Primer-BLAST^59^ under default parameters with the requirement for exon-exon junction spanning specified. qPCR was performed using Power SYBR Green Master Mix (Applied Biosystems). Reactions were run in triplicate, following default SYBR Green Standard cycle specifications on the Viia7 Real-Time PCR System (Applied Biosystems). Relative quantification followed the 2^-ΔΔCT^ method, normalizing results to *Actb* in the EGFP-aliquot of cells for each region, respectively.

### Correlation analysis between regions and within replicates

Peaks from all six ATAC-seq libraries and the two “Joint” ATAC-seq libraries were concatenated together, sorted on the basis of chromosomal location, merged into a unified peak set^60^, and converted to Simplified Annotation Format (SAF). Reads from each BAM file overlapping this unified peak set were quantified with the Rsubread package “featureCounts” command, with the following options: isPaired End = TRUE, requireBothEndsMapped = FALSE. Read counts were normalized for each library using conditional quantile normalization^61^, accounting for library size, peak length, and peak GC content. Pearson correlation co-efficients were calculated from this normalized count matrix and visualized using corrplot^62^ and RColorBrewer^63^ and LSD^64^.

### Sequence constraint analysis

Average phastCons^65^ were calculated for the “Joint” peak file for both the MB and FB libraries using Cistrome^66^ (http://cistrome.org/ap/). Beforehand, peaks with overlap of exons or promoters (defined here as +/-2000bp from the transcriptional start site) were removed. The exon and promoter BED files were downloaded from the UCSC table browser^67^ (Mouse genome; mm9 assembly; Genes and Gene Predictions; RefSeq Genes track using the table refGene).

### Gene ontology of nearest expressed gene

The Genomic Regions Enrichment of Annotations Tool^68^ (GREAT; v3.0.0; http://great.stanford.edu) predicted the GO term enrichment in the catalogues. Beforehand, peaks were processed to: a) remove peaks overlapping commonly open regions; b) select the top 20,000 peaks and; c) overlap the nearest expressed gene’s transcriptional start site (TSS).

First, regions that are commonly open were defined as those regions of the genome that are open in >30% of ENCODE DNase hypersensitivity site (DHS) assays in mouse tissues. These ubiquitously open regions were removed from the peak files. Next, to limit the number of regions submitted to GREAT such that the binomial distribution for calculating fold enrichment values was still valid, peak files were limited to the top 20,000 peaks on the basis of q-value.

Finally, in order to limit ourselves to the nearest expressed gene, we supplied a list of the TSSs of the nearest expressed gene that are in the GREAT database. The list of genes and their TSSs used by GREAT was downloaded from: http://bejerano.stanford.edu/help/download/attachments/2752609/mm9.great3.0.genes.txt. Only genes that are in this list with RPKM > 1 were considered as expressed. The nearest expressed gene to each of the top 20,000 peaks was identified. Each peak is associated with its nearest expressed gene and to ensure that GREAT only considered these nearest genes for analysis, we submitted these nearest expressed gene’s TSSs as a proxy for each peak. These proxy peaks were submitted to GREAT using the NCBI build 37 (mm9) assembly, under whole genome background regions, with the single nearest gene as the association rule, including curated regulatory domains.

### Quantification of overlap between CRE catalogues and the VISTA Enhancer Browser

All elements tested *in vivo* were downloaded from the VISTA Enhancer Browser (https://enhancer.lbl.gov) on September 4, 2016. These regions were stratified into those annotated as positive or negative. BED co-ordinates of these regions were extracted and intersected with the ATAC-seq catalogues. Positive regions were further stratified into those with annotations for only forebrain, only midbrain, only hindbrain, combinations of regions (“Multiple regions”), all three regions (“Whole brain”), summing to the “Neuronal” category, or were annotated as positive but driving expression in none of those three regions (“Non-neuronal”).

### Testing five putative CREs for *in vivo* reporter activity

Prioritized regions were PCR amplified (**Supplementary Table 5**) from human gDNA and cloned into either pENTR for mouse *lacZ* assays (Invitrogen) or pDONR221 for zebrafish assays (Invitrogen). Regions were sequence validated and LR cloned (Invitrogen) into either an *hsp68-lacZ* vector or pXIG vector, with a TdTomato cassette in place of GFP.

Generation of transgenic mice and E11.5 embryo staining was performed as previously described^69–71^ using FVB strain mice. Embryos expressing the *lacZ* reporter gene were scored and annotated for their expression patterns by multiple curators. For a construct to be considered positive, a minimum of three embryos per construct were required to demonstrate reporter activity in the same tissue. Mouse transient transgenic assays were approved by the Lawrence Berkeley National Laboratory Animal Welfare and Research Committee.

Generation of transgenic zebrafish was performed as previously described^72^ in AB zebrafish. At 3dpf and 5dpf, reporter expression patterns were evaluated. For a construct to be considered as positive, ≥25% of mosaic embryos had to display reporter activity in one or more anatomical structures. Positive zebrafish were quantified for reporter activity in five anatomical regions (forebrain, midbrain, hindbrain, amacrine cells, spinal cord).

### Regulatory vocabulary development

We applied the machine learning algorithm gkm-SVM to our MB and FB catalogues, under default settings. We trained on the sequences underlying the summits ±250bp of non-ubiquitously open, top 10,000 peaks by signal intensity, versus five negative sets, matched for GC content, length, and repeat content. Weights across all five tests were averaged for all 10-mers.

All 10-mers with weight ≥1.50 were clustered on sequence similarity using Starcode^73^, using sphere clustering with distance set to 3. clustalOmega^74^ aligned the sequences within these clusters and MEME^75^, under default parameters, excepting ‐dna ‐maxw 12, generated position weight matrices of these aligned clusters. Tomtom^76^, querying the Jolma 2013, JASPAR Core 2014, and Uniprobe mouse databases, identified the top transcription factors corresponding to these position weight matrices, under default parameters excepting ‐no-ssc ‐min-overlap 5 ‐evalue ‐thresh 10.0.

The same procedure was used to identify transcription factors specifically conveying regulatory potential in the MB library relative to the FB library, except during gkm-SVM training, the positive set was specified as the top 10,000 non-ubiquitously open MB summits and the negative set was specified to be the top 10,000 non-ubiquitously open FB summits, both ±250bp.

### Transcription factor footprinting

CENTIPEDE^77^ was used to identify footprints. Sequences underlying the deeply sequenced MB library peaks, less those ubiquitously open, were extracted. FIMO^78^, with options ‐‐text ‐‐parse-genomic-coord, identified all locations underlying ATAC-seq peaks of the motifs identified above. Additionally, conservation data from 30-way vertebrate phastCons was considered in the CENTIPEDE calculations; for each PWM site, those with mean conservation score greater than 0.9 were considered. Finally, the BAM file read end co-ordinates were adjusted in response to the shift in co-ordinates due to the transposase insertion^79^. As such, following the original ATAC-seq method^17^, reads were adjusted +4bp on the positive strand and ‐5bp on the negative strand.

### Genome-wide read pileup over predicted motif sites

FIMO, as above, was used to identify all co-ordinates genome wide of the identified motifs. deepTools^80^ “bamCoverage” tool was run under default conditions, to convert the deeply sequenced MB library BAM to bigwig format. Following this, a matrix file was generated with “computeMatrix”, with options ‐‐ referencePoint center ‐b 1000 ‐a 1000 ‐bs 50 specified. Finally, "plotHeatmap" was used to generate plots indicating ATAC-seq read pileup over predicted motif sites.

### *In vivo* validation of the MB-specific enhancer

The MB-specific peak was PCR amplified (**Supplementary Table 5**) from human genomic DNA and TA cloned into pCR8 (Invitrogen). Regions were sequence validated and LR cloned (Invitrogen) into either an *hsp68-lacZ* vector or a modified pXIG vector, with a TdTomato cassette in place of GFP.

For zebrafish transgenesis, the modified pXIG vector was injected into 1-2 cell stage embryos as previously described^72^ in AB zebrafish. TdTomato reporter expression was assayed at 72hpf and 5dpf; mosaic embryos positive for TdTomato expression were selected and raised to adulthood and founders were identified. Progeny of founders were screened at 72hpf for reporter activity.

For mouse transgenesis, the generated *hsp68-lacZ* vector was purified in a double CsCl gradient (Lofstrand Labs Ltd) and stable mouse transgenesis was performed in C57BL/6 mice by Cyagen Biosciences Inc. Multiple founder lines were generated. For lacZ staining, embryos were collected at E12.5, and mouse brains were isolated at E15.5, P7, P30, and P574. Brains were roughly sectioned in 1mm sections at P7 and P30 and animals were perfused at P574 and fixed brains were sectioned (200μm) with a vibratome. Specimens were subsequently fixed for 2 hours on ice in 1% formaldehyde, 0.2% glutaraldehyde, 0.02% Igepal CA-630 in PBS. Following fixation, tissues were permeabilized over 3×15 minute washes in 2mM MgCl2 and 0.02% Igepal CA-630 in PBS at room temperature. Embryos/tissues were incubated overnight at 37°C in staining solution, containing 320μg/mL X-Gal in N,N-dimethyl formamide, 12mM K-ferricyanide, 12mM K-ferrocyanide, 0.002% Igepal CA-630, 4mM MgCl_2_ in PBS. Specimens were washed in 0.2% Igepal CA-630 in PBS over 2×30 minutes and finally stored in 4% formaldehyde, 100mM sodium phosphate, and 10% methanol.

### Patient sequencing and genotyping at *SNCA*

A total of 986 PD patients and 992 controls who were seen at the Mayo Clinic in Jacksonville, FL were sequenced across the putative enhancer and genotyped for 25 variants across the *SNCA* locus. The variants chosen for genotyping were confirming those identified by sequencing of the enhancer as well as assessing those identified in Guella *et al*^41^. For PD patients, median age at blood draw was 69 years (Range: 28-97 years), median age at PD onset was 67 years (Range: 28-97 years), and 631 patients (64.0%) were male. Median age at blood draw in controls was 67 years (Range: 18-92 years) and 415 subjects (41.8%) were male. Patients were diagnosed with PD using standard clinical criteria^81^. All subjects are unrelated non-Hispanic Caucasians of European descent. The Mayo Clinic Institutional Review Board approved the study and all subjects provided written informed consent.

Genomic DNA was extracted from whole blood using the Autogen FlexStar. Sanger sequencing of the enhancer region was performed bidirectionally using the ABI 3730xl DNA analyzer (Applied Biosystems) standard protocol. Sequence data was analyzed using SeqScape v2.5 (Applied Biosystems). Statistical analyses were performed using both SAS and R^82^. Of the variants identified within the enhancer, only those with minor allele frequency greater than 5% were evaluated for association with PD in single-variant analysis. Associations between individual variants and PD were evaluated using logistic regression models, adjusted for age at blood draw and sex, and where variants were considered, under an additive model (i.e. effect of each additional minor allele). Odds ratios and 95% confidence intervals were estimated and a Bonferroni correction for multiple testing, due to the four common variants that were evaluated for association with PD, was utilized in single-variant analysis, after which p-values ≤ 0.0125 were considered as statistically significant.

Genotyping the 25 SNPs across the SNCA locus was performed using the iPLEX Gold protocol on the MassARRAY System and analysed with TYPER 4.0 software (Agena Bioscience). For the 25 SNPs genotyped across the *SNCA* locus, all genotype call rates were >95% and there was no evidence for departure from Hardy-Weinberg equilibrium (all *χ*^2^ p-values > 0.05 after Bonferroni correction). Haplotype frequencies in cases and controls was estimated using the haplo.stats package^83^ function “haplo.group”. Associations between haplotypes and risk of PD were evaluated using score tests of association^84^ using the “haplo.score” function. Tests were adjusted for age at blood draw and sex, haplotypes occurring in less than 1% of subjects were excluded, and only individuals with no missing genotype calls for any variants were included. A Bonferroni correction for multiple testing was applied, after which p-values ≤ 0.0042 were considered as statistically significant, due to the 12 different common haplotypes that were observed and tested for association with PD risk.

### Protein array testing differential binding

HuProt v3.1 human proteome microarrays printed on the PATH surface containing >20,000 unique proteins representing 16,152 genes (CDI laboratories)^40^ were blocked with 25mM HEPES pH 8.0, 50mM K Glutamate, 8mM MgCl_2_, 3mM DTT, 10% glycerol, 0.1% Triton X-100, 3% BSA on an orbital shaker at 4^°^C for ≥3 hours. Allele specific protein-DNA binding interactions were identified through dye-swap competition of major and minor alleles labeled with either Cy3 or Cy5. DNA fragments for rs2737024 and rs2583959 were synthesized with the SNP for each allele flanked by 15 nucleotides of the upstream and downstream sequence and a common priming site at the 3’ end (**Supplementary Table 5**).

The dsDNA fragments were created by separately annealing a primer containing a Cy3 or Cy5 label and adding Klenow (NEB) with dNTP to fill-in the complementary strand for each allele^85^. Cy3 labeled major allele was mixed with Cy5 labeled minor allele (each at 40nM) in 1x hybridization buffer (10mM TrisCl pH 8, 50mM KCl, 1mM MgCl_2_, 1mM DTT, 5% glycerol, 10μM ZnCl_2_, 3mg/mL BSA) and added to an array, dyes were then swapped for each allele and the mixture was then added to a second array. DNA was allowed to bind overnight at 4^°^C on an orbital shaker with protection from light. Chips were washed once with cold 1xTBST (0.1% Triton X-100) for 5 minutes at 4^°^C, rinsed, and dried in the centrifuge. Cy5 and Cy3 images were taken separately on a Genepix 4000B scanner and, after alignment to the ․gal file, individual spot intensities were extracted using the Genepix Pro software.

Allele specific interactions were identified through dye swap analysis. The ratio of major/minor allele binding was calculated using the duplicate spot average median foreground signal for each protein according to the following equation: 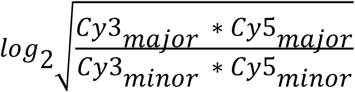

Mean intensity was calculated by averaging the foreground signal for the Cy3 and Cy5 channels of the major and minor alleles. MA plots were made for each allele using the calculated mean intensity and the log ratio of the major/minor allele.

## DATA AVAILABILITY

ATAC-sequencing and RNA-sequencing data will be available at the Gene Expression Omnibus (GEO) under the accession number GSEXXXXXX.

## ACKNOWLEDGEMENTS

This research undertaken at Johns Hopkins University School of Medicine was supported in part by awards from NIH (NS62972 and MH106522 to ASM; GM111514 to HZ; HG007348 to MAB). The Mayo Clinic collection was supported in part by a Morris K. Udall Center of Excellence in Parkinson’s disease Research (P50 NS072187), American Parkinson’s Disease Association Center and The Mangurian Foundation for Lewy body research. OAR is supported by NS078086 and NS10069 (NIH), W81XWH-17-1-0249 (Department of Defense), The Michael J. Fox Foundation and The Little Family Foundation. ZKW is supported by the Mayo Clinic Center for Regenerative Medicine, Mayo Clinic Center for Individualized Medicine, Mayo Clinic Neuroscience Focused Research Team (Cecilia and Dan Carmichael Family Foundation, and the James C. and Sarah K. Kennedy Fund for Neurodegenerative Disease Research at Mayo Clinic in Florida), the gift from Carl Edward Bolch, Jr., and Susan Bass Bolch, The Sol Goldman Charitable Trust, and Donald G. and Jodi P. Heeringa. Research conducted at the E.O. Lawrence Berkeley National Laboratory was performed under U.S. Department of Energy contract DE-AC02-05CH11231, University of California and was supported by HG003988 (NIH) to LAP.

## AUTHOR CONTRIBUTIONS

SAM, ASM designed the study and wrote the paper. SAM, PWH, XR, WDL, SJK, and ELW performed various experiments. Transgenic experiments were performed by SAM, NJB, and JFT (zebrafish) and by SAM, JAA, DED, AV, and LP (mice). SAM and MAB performed the gkm-SVM analyses. PD patient sequencing and analysis was performed by SAM, AIS, MGH, NND, ZKW, and OAR. SAM, CDM, and HZ performed and analysed differential protein array binding assays. SAM implemented the computational algorithms to process the raw data and conduct analyses thereof. SAM and ASM analyzed and interpreted the resulting data. SAM contributed novel computational pipeline development. SAM and ASM wrote the manuscript, with all other authors contributing. Correspondence to ASM.

